# DAGBagM: Learning directed acyclic graphs of mixed variables with an application to identify prognostic protein biomarkers in ovarian cancer

**DOI:** 10.1101/2020.10.26.349076

**Authors:** Shrabanti Chowdhury, Ru Wang, Qing Yu, Catherine J. Huntoon, Larry M. Karnitz, Scott H. Kaufmann, Steven P. Gygi, Michael J. Birrer, Amanda G. Paulovich, Jie Peng, Pei Wang

## Abstract

**Motivation:** Directed gene/protein regulatory networks inferred by applying directed acyclic graph (DAG) models to proteogenomic data has been shown effective for detecting causal biomarkers of clinical outcomes. However, there remain unsolved challenges in DAG learning to jointly model clinical outcome variables, which often take binary values, and biomarker measurements, which usually are continuous variables. Therefore, in this paper, we propose a new tool, DAGBagM, to learn DAGs with both continuous and binary nodes. By using appropriate models for continuous and binary variables, DAGBagM allows for either type of nodes to be parents or children nodes in the learned graph. DAGBagM also employs a bootstrap aggregating strategy to reduce false positives and achieve better estimation accuracy. Moreover, the aggregation procedure provides a flexible framework to robustly incorporate prior information on edges for DAG reconstruction.

**Results:** As shown by simulation studies, DAGBagM performs better in identifying edges between continuous and binary nodes, as compared to commonly used strategies of either treating binary variables as continuous or discretizing continuous variables. Moreover, DAGBagM outperforms several popular DAG structure learning algorithms including the score-based hill climbing (HC) algorithm, constraint-based PC-algorithm (PC-alg), and the hybrid method max-min hill climbing (MMHC) even for constructing DAG with only continuous nodes. The HC implementation in the R package DAGBagM is much faster than that in a widely used DAG learning R package bnlearn. When applying DAGBagM to proteomics datasets from ovarian cancer studies, we identify potential prognostic protein biomarkers in ovarian cancer.

**Availability and implementation:** `DAGBagM` is made available as a github repository https://github.com/jie108/dagbagM.

## 1 Introduction

Ovarian cancer is the most lethal gynecological malignancy and is often diagnosed at an advanced stage (Huang et al., 2010). The daunting five-year survival rate of ovarian cancer patients remains largely unchanged during the past few decades, despite all the efforts and resources devoted to genomic research of this disease. Therefore, there is an urgent need to develop novel approaches to identify new biomarkers/targets for diagnosis, prognosis or treatment of ovarian cancer patients.

Recent breakthrough in proteomics research has made it possible to monitor tens of thousands of proteins in one biological sample simultaneously. High throughput proteomics experiments have been performed on ovarian tumor samples by the *Clinical Proteomic Tumor Analysis Consortium* (*CPTAC*) (Zhang et al., 2016; McDermott et al., 2020; Arshad et al., 2019), which provides an unprecedented opportunity to screen for potential prognostic protein biomarkers that may not otherwise be discovered using previously defined genomic approaches due to the large amount of post translational modifications in cells. In addition, like most other cancers, ovarian cancer is a complex disease, involving complicated pathway interactions and dysfunctions across multiple biological processes. Thus, a systems-level approach is the key for enhancing our understanding of molecular mechanism underlying the disease, and for detecting effective biomarkers for cancer progression and metastasis. Consequently, higher order molecular networks could serve as central tools for extracting relevant information from high dimensional proteogenomic data. Therefore, in this paper, we aim to screen for prognostic protein biomarkers, which have causal associations with ovarian cancer prognosis, via constructing *directed acyclic graphs* (*DAG*) based on CPTAC proteomics data.

In protein regulatory networks, edges are mathematical representations that characterize the relationships among proteins in the cell. These relationships are further used to understand the detailed steps for the formation of protein complexes or the signaling pathways relating to the drug targets. Often the goal of such network construction is to identify modules or clusters containing regulators of major drivers that show causal associations with clinical outcomes. Among different approaches of constructing regulatory networks, DAG models are commonly used to infer causality of the regulatory relationships among the interacting entities (e.g., genes or proteins) in a complex biological system (Pearl, 2000; Zhu et al., 2012; Sung et al., 2016; Sung and Hua, 2016; Friedman et al., 2000; Peer et al., 2001; Sachs et al., 2005). DAG models are also often used in prognostic studies to identify causal associations between biomarkers and clinical variables (Williams et al., 2018; Asvatourian et al., 2018). Besides constructing gene/protein regulatory networks, DAG models are also used in other applications such as natural languages processing (Bishop et al., 2006), developing medical intelligence systems, expression quantitative trait loci mapping (Neto et al., 2010), among others.

Despite considerable efforts and many pioneering works, there remains challenges in DAG structure learning, especially when the node set contains both continuous and discrete variables (i.e., mixture nodes). While discretization of continuous nodes is a commonly used strategy (Scutari, 2009), it does not always guarantee the preservation of the original dependence structure and may also lead to loss of information. On the other hand, simply treating discrete variables as continuous variables leads to model misspecification and false edges/directions. However, in practice, the need to deal with mixed nodes are very common. For example, the clinical outcomes are often binary end-points, e.g, patient response to treatment or survival status, whereas potential biomarkers are often continuous measures such as protein expression levels. In (Andrews et al., 2018), two scoring methods are proposed to handle mixed nodes: One is based on conditional Gaussian distributions and thus is biased towards having discrete parents of continuous children, and the other is based on higher order polynomial approximation which mainly aims at modeling nonlinear relationships among the nodes. In (Zhu et al., 2012), the authors, who aimed to construct DAGs for discrete genetic variables and continuous functional molecular variables, also constrained that discrete variables can only serve as the parent nodes but not the child nodes.

In this paper, aiming to search for biomarkers causally associated with clinical variables of interest, we propose a score based DAG structure learning algorithm, `DAGBagM`, that models the continuous nodes by conditional Gaussian distributions and the binary nodes through logistic regressions. Particularly, DAGBagM allows for both continuous and binary variables to be children nodes and is not biased towards a particular type of edges. Moreover, to tackle computational challenges associated with large number of nodes, we develop an efficient implementation of the *hill climbing algorithm* where at each search step information from the previous step is utilized to speed up both score calculation and acyclic status check.

DAG structure learning also tends to be highly variable even with moderate number of nodes. The learned graph tends to change drastically with even small perturbation of the data. To tackle this challenge, DAGBagM employs a novel aggregation procedure inspired by *bootstrap aggregating* (*bagging*) (Breiman, 1996) and couples this procedure with the aforementioned score-based algorithm. Data perturbation and model aggregation have been previously considered for DAG learning ((Friedman et al., 1999; Imoto et al., 2002; Elidan, 2011; Elidan et al., 2002; Broom et al., 2012)). In `DAGBagM`, through a distance metric defined on the DAG space, our aggregation procedure results in a valid DAG based on an ensemble of DAGs learned on bootstrap resamples of the data. As shown by simulation studies, this aggregation strategy is able to greatly reduce false positives with only moderate sacrifice in power.

`DAGBagM` is also flexible in taking into prior information of edge directions. Prior information can be important for DAG structure learning as edge directions are not always identifiable without external information. Independent resources such as time course experiments could provide valuable information on regulatory directions. Thus, it is helpful to incorporate prior knowledge/information from independent sources when performing DAG learning. On the other hand, often all prior knowledge/information is not accurate. Therefore, we utilize priors through the aggregation process to enhance robustness. Specifically, in each bootstrap iteration, `DAGBagM` not only sample the data but also the priors, such that false priors shall only impact a subset of the models in the ensemble.

In the real data application, we perform DAG analysis to screen for proteins causally associated with tumor prognosis in ovarian cancer, focusing on the metabolic pathways such as Oxidative Phosphorylation and Adipogenesis, as studies have shown that metabolic reprogramming in ovarian cancer are key factors underlying the metastasis of cancer cells (Han et al., 2018). We implemented an integrative learning pipeline using `DAGBagM` to borrow information across multiple proteomics data sets of ovarian cancer and identified multiple novel prognostic proteins. The results shed light on the underlying role of the key markers of the metabolic pathways that are causally associated with cancer prognosis and hence can potentially contribute to overcome the non-response towards anti-cancer agents.

The rest of the paper is organized as follows. In Section 2 we provide a detailed description of the proposed `DAGBagM` algorithm. The first part of Section 3 presents numerical results from simulation studies; and the second part details the application on proeteomics ovarian cancer datasets. We conclude the paper with a discussion and further details can be found in the supplementary material.

## 2 Method: DAGBagM

In this section we present a new tool – `DAGBagM` – for learning directed acyclic graphs with both continuous and binary nodes. In `DAGBagM`, we adopt a score-based approach coupled with bootstrap aggregation. The score calculation uses separate distributions for continuous and binary nodes which performs better than the strategies of either treating all nodes as continuous or discretizing continuous nodes. Furthermore, the bootstrap aggregation step greatly reduces the number of false positives and improves on reproducibility of the results. We also have a very efficient implementation of the hill climbing search algorithm which enables applying `DAGBagM` to learn graphs with a large number of nodes.

A directed acyclic graph 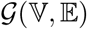 consists of a node set 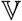 and an edge set 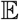 with directed edges from *parents* nodes to *children* nodes. In a DAG model, the node set corresponds to a set of random variables and the edge set encodes the conditional dependence relationships among these random variables. DAG structure learning amounts to identifying the parent(s) set (also referred to as neighborhood) of each node in the graph. Although different DAGs could encode the same set of conditional dependencies (which form an equivalent class of DAGs), it is shown that, two DAGs are equivalent if and only if they have the same set of skeleton edges and *v*—structures ((Verma and Pearl, 1991)). The skeleton edges are obtained by removing directions from the directed edges and a *v*—structure is a triplet of nodes (*x*_1_, *x*_3_, *x*_2_), such that 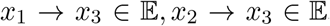, and *x*_1_, *x*_2_ are not adjacent. There are mainly three classes of methods for DAG structure learning, namely, *score-based* methods (e.g., (Geiger and Heckerman, 1994)), *constraint-based* methods, e.g., PC algorithm (`PC-alg`) (Verma and Pearl, 1991; Spirtes et al., 2001; Kalisch and Bühlmann, 2007) and *hybrid* methods, e.g., Max-Min Hill Climbing (MMHC) (Tsamardinos et al., 2006).

In score-based methods, DAG structure learning is treated as a model selection problem through minimizing a pre-specified *score* (e.g., a penalized negative log likelihood score) over the space of DAGs defined on a given set of nodes. When all nodes are continuous variables, they are often modeled as jointly Normally distributed. On the other hand, if all nodes are discrete, multinomial distributions may be used. When there are both continuous nodes and discrete nodes, a common practice is to discretize the continuous nodes. However, this does not guarantee the preservation of the dependence structure and it may also lead to loss of power. Alternatively, one may treat the discrete nodes as continuous, which however leads to model misspecification and false edges/ directions. In `DAGBagM`, we treat continuous nodes and binary nodes separately. Specifically, for continuous nodes, we model them as normally distributed given their parent(s) and for binary nodes we model them through logistic regressions.

One challenge of score-based methods is that due to the super-exponentially large DAG space, an exhaustive search for optimal models is usually computationally infeasible. Therefore, greedy search algorithms are often employed. One of the most popular search algorithms is the *hill climbing* (*HC*) *algorithm* (Russell et al., 2010), which performs local search at each step for the best operation that results in the maximum score improvement among a set of eligible operations at the current step. The graph will be updated according to the selected best operation and the search will stop if no improvement is found. The major computational cost of the HC algorithm comes from the score calculation and acyclic status check which are needed for each eligible operation at each search step.

We develop an efficient implementation of the HC algorithm that uses information from previous steps to facilitate the score calculation and acyclic status check in the current step. Details are given in two propositions in A.1 of the Supplementary Material. The key observation is that for majority of the operations, score change and acyclic status remain the same as those in the previous step and thus do not need to be re-assessed. E.g., any operation that does not involve the neighborhood(s) changed by the selected operation from the previous step, results in the same score change as in the previous step, and hence re-calculation is not needed.

`DAGBagM` also employs a novel aggregation procedure to learn stable structures and to reduce false positive edges. We use structural hamming distance, commonly used in information theory and in evaluation of DAG learning results (Tsamardinos et al., 2006; Perrier et al., 2008), to define a distance metric on the DAG space (of a given set of nodes). We first learn an ensemble of DAGs where each DAG is learned on a bootstrap resample of the data (by applying the proposed score-based method). We then obtain an aggregated DAG through minimizing the average distance to DAGs in the ensemble (by applying the HC algorithm). Moreover, `DAGBagM` can incorporate prior information through blacklist(s) of forbidden edges and/or whitelist(s) of edges that should always be in the graph. This is done through the exclusion of certain operations from the set of eligible operations at each updating step. Blacklist or whitelist may be utilized in either the individual DAG learning step or the aggregation step.

Major steps of `DAGBagM` are summarized in Figure 1 and in Algorithm 1. More detailed information regarding each step is provided in subsequent subsections. Note that, although we describe the `DAGBagM` algorithm under the situation when there are both continuous and binary nodes, it is applicable when there are only continuous nodes or when there are only binary nodes. Moreover, both continuous nodes and binary nodes could be either parent nodes or child nodes.

**Figure 1:**
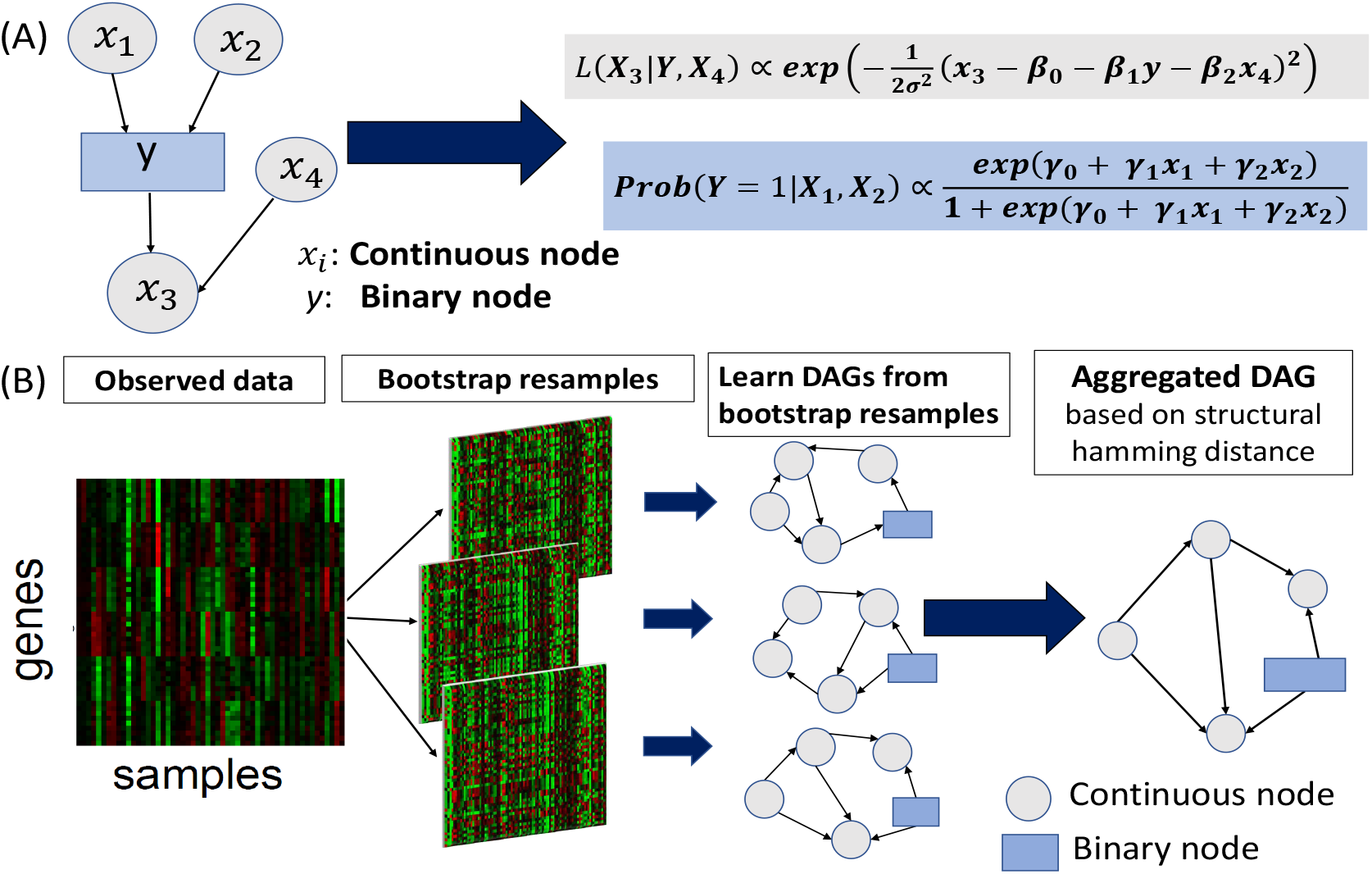
DAGBagM algorithm. (A) shows a DAG with continuous nodes modeled by conditional Gaussian distributions and a binary node modeled through logistic regressions. (B) shows major steps of the `DAGBagM` algorithm.

### 2.1 Score calculation

Structure learning based on the likelihood score overfits the data since it always favors larger models, i.e., distributions with less independence constraints/DAGs with more edges. Therefore, it is reasonable to consider scores that penalize for model complexity. Since Bayesian information criterion (BIC) (Schwarz, 1978) is *model selection consistent* and *locally consistent* (Chickering, 2002), `DAGBagM` adopts BIC-type scores to be minimized by the hill-climbing algorithm.

For a continuous node, denoted by *X*, at each search step, its score is calculated by regressing *X* onto its current parent(s). Specifically, for a given graph 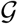, 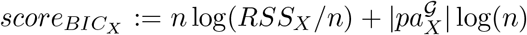, where, *RSS_X_* is the residual sum of squares, *n* is the sample size and 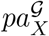 denotes the parent set of *X* in graph 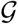. For a binary node *Y*, the score is obtained by regressing *Y* onto its current parent set through logistic regression:

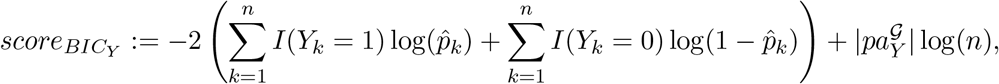

where 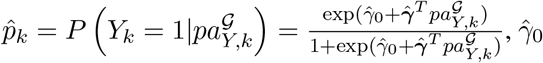 is the fitted intercept and 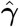 is the vector of the fitted coefficients in logistic regression. Finally, the score of a graph 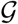 is the summation of individual node scores: 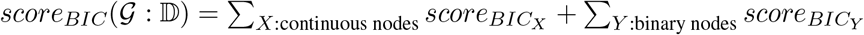, where 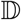 denotes the data.

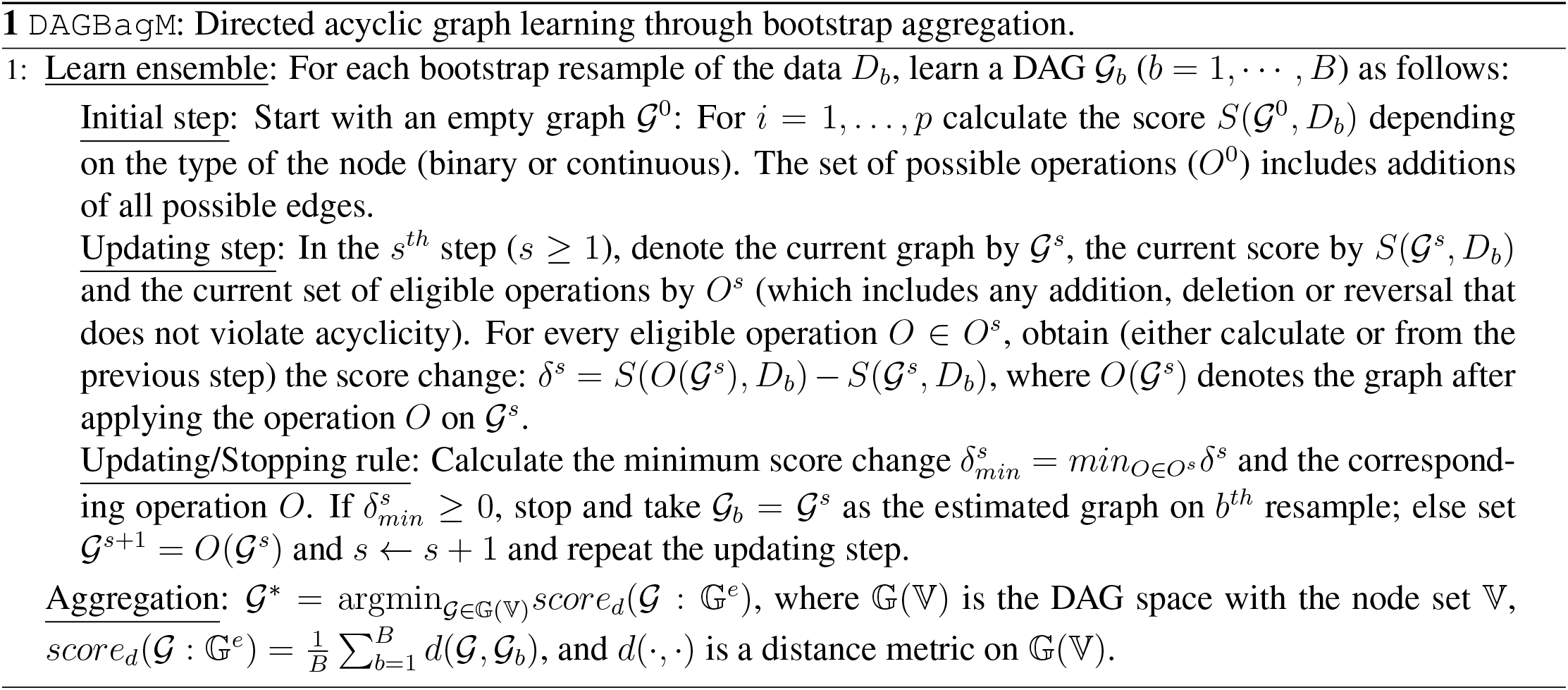

### 2.2 Bootstrap aggregation

The aggregation of an ensemble of DAGs is nontrivial because the notion of mean is not straightforward on the DAG space. Here, we generalize the idea of mean by searching for a DAG that minimizes an average distance to the DAGs in the ensemble. For this purpose, we define a distance metric based on the *Hamming distance*. In information theory, the Hamming distance between two 0-1 vectors of equal length is the minimum number of substitutions needed to convert one vector to another. This can be generalized to give a distance measure between two DAGs with the same set of nodes, defined as the minimum number of basic operations, namely, addition, deletion and (possibly) reversal that are needed to convert one graph to another. This definition leads to a valid distance metric and is referred to as the *structural Hamming distance* (*SHD*).

While there are different variants of SHD depending on how the reversal operations are counted, here we focus on the case where the reversal operation is counted as one unit of operation. This leads to the following distance: 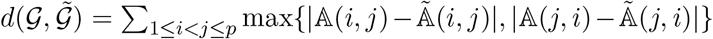, where 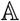 and 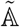 denote the adjacency matrices of the DAGs 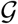 and 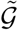, respectively. The adjacency matrix of a DAG is a 0-1 element matrix where the (*i, j*)-th element is one if there is a directed edge from the *i*th node to the *j*th node; otherwise it is zero.

Finally the *aggregation score* between a DAG 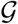 and an ensemble of DAGs 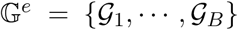 is the average distance between 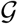 and the DAGs in the ensemble: 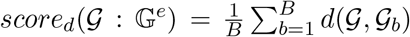. By Proposition 3 in A.2 of the Supplementary Material, the aggregation score 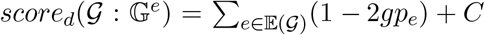, where *C* is a constant, and *gp_e_* is a generalized selection frequency. Given an ensemble, one can search for the DAG that minimizes the aggregation score while maintaining acyclicity by applying the HC algorithm. We defer details to the Supplementary Material.

## 3 Results

### 3.1 Simulation studies

In this section, we conduct simulation studies to examine the proposed `DAGBagM` algorithm and compare it to several existing DAG structure learning algorithms.

#### Simulation Setup

We perform two sets of simulation experiments with (i) only continuous nodes; and (ii) both continuous and a single binary node.

For scenario (i), given the true data generating graph 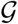 and sample size *n*, i.i.d. samples are generated according to *Gaussian linear mechanisms*: 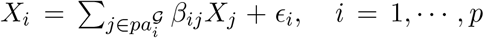, where *ϵ_i_*’s are independent Gaussian random variables with mean zero and variance 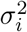. The coefficients *β_ij_* s are uniformly generated from 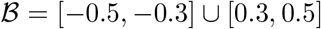 and the error variances 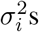 are chosen such that for each node the corresponding *signal-to-noise-ratio* (*SNR*), defined as the ratio between the standard deviation of the signal part and that of the noise part, is within [0.5, 1.5]. For the purpose of comparison, we also consider the non-aggregated hill climbing algorithm (`HC`) (our own implementation), the constraint-based algorithm `PC-alg` (implemented in R package *pcalg* (Kalisch et al., 2012) and use α = 0.005 as suggested in (Kalisch and Bühlmann, 2007)), and the hybrid algorithm MMHC as implemented in R package *bnlearn* (Scutari, 2010) and use the default tuning parameter at 0.05.

For scenario (ii), we use a similar data generating scheme for the continuous nodes as in (i). The binary node, denoted by *Y*, is generated by a logistic regression model: 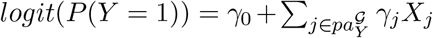. We consider two methods for comparison, namely `DAGBagM`_*C*_ and `bnlearnD`. In `DAGBagM`_*C*_, we simply apply the `DAGBagM` algorithm while treating all nodes (including the binary node) as continuous. In `bnlearnD`, we first discretize every continuous node using the median cut-off. We then treat all nodes as binary nodes and apply the “hc” function with *BIC* score implemented in the R package *bnlearn* (Scutari, 2010). For a fair comparison with `DAGBagM` based methods, we do this on each bootstrap resample and then apply the proposed aggregation algorithm to learn an aggregated DAG.

For each simulation setting, 100 independent replicates are generated. We consider DAGs with different numbers of nodes *p* as well as different sample sizes *n*. The true DAGs are shown in Figures A.1-A.2 of Section A.4 in the Supplementary Material.

#### Simulation results

We report results on power and false discovery rate (FDR) in terms of detection of skeleton edges (i.e., without direction) in tables below. Power is calculated as the ratio between the number of correct edges in the estimated DAG to the number of total edges in the true DAG, and FDR is calculated as the ratio between the number of false edges in the estimated DAG to the number of total edges in the estimated DAG. All numbers are averaged over results on 100 replicates. Along with power and FDR, we also report the F1 score defined as 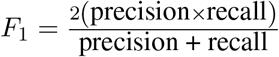, where precision=1-FDR and recall=power.

For scenario (i) – only continuous nodes, we first consider an empty graph with *p* = 1000 nodes, 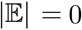 edge and sample size *n* = 250 to illustrate the effect of aggregation in reducing the number of false positive edge detections. Note here any detected edge would be a false positive. As can be seen from Table 1, `DAGBagM` results in very few false positives, whereas the three non-aggregation methods, namely, `HC, PC–alg` and `MMHC`, all have large number of false positives.

**Table 1:** **Simulation scenario (i)** – only continuous nodes. True DAG: *p* = 1000 nodes, 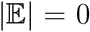 edge; Sample size *n* = 250

| Method | Total # of detected edges | Total # of detected v-structures |
| --- | --- | --- |
| DAGBagM | 5.7 (2.83) | 0 (0) |
| HC | 1995.6(2.12) | 3068.3(124.53) |
| PC-alg | 946.5 (12.47) | 560.8 (22.44) |
| MMHC | 4199.6 (45.7) | 9821.1 (248.11) |

We then consider a graph (Supplementary Figure A.1) with *p* = 504 nodes, 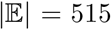 edges under two sample sizes, *n* = 100 and *n* = 250. It is clear from Table 2 that `DAGBagM` outperforms the other three methods in terms of balancing power and FDR as measured by the *F*_1_ score. It is also obvious that the larger sample size leads to better performance for all methods, especially so for `DAGBagM`.

**Table 2:** **Simulation scenario (i)** – only continuous nodes. True DAG: *p* = 504 nodes, 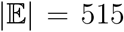 edges; Sample size *n* = 100, 250, SNR ∈ [0.5, 1.5]

| Method | $n = 100$ | | | $n = 250$ | | |
| --- | --- | --- | --- | --- | --- | --- |
| | power | FDR | $F_1$ score | power | FDR | $F_1$ score |
| DAGBagM | 0.66 | 0.02 | 0.79 | 0.9 | 0.02 | 0.94 |
| HC | 0.84 | 0.57 | 0.57 | 0.96 | 0.49 | 0.67 |
| PC-alg | 0.61 | 0.68 | 0.42 | 0.79 | 0.08 | 0.85 |
| MMHC | 0.72 | 0.67 | 0.45 | 0.8 | 0.29 | 0.75 |

For scenario (ii) – mixture of continuous nodes and one binary node, we consider graphs (Supplementary Figure A.2) with different combinations of number of nodes and edges. In each graph, there is a single binary node which is the child of two continuous parents, and is the parent of one continuous child. These settings mimic the third step in the ovarian cancer application where we learn DAGs on modules containing 10 to 20 continuous biomarkers and one binary clinical outcome. Note that although here we focus on one binary node to mimic real data application, `DAGBagM` can handle any number of continuous and binary nodes.

In Table 3 we report edge detection results under varying sample sizes for a graph (Supplementary Figure A.2 A) with *p* = 11 nodes and 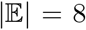 edges. In Table 4, we report edge detection results for graphs with different combinations of *p* and 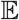 (Supplementary Figure A.2 B - E) for a fixed sample size *n* = 100. As can be seen from the two tables, the performances of `DAGBagM` and `DAGBagM`_*C*_ are quite similar in terms of power and FDR of skeleton edges detection, while the performance of `bnlearnD` is much worse under all cases. Quite obviously, the performance of all three methods improves with the increase of sample size *n* (Table 3), while with the increase of *p* and 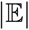, the performance becomes worse for all three methods (Table 4). In both Table 3 and Table 4 we also report the percentage of edge detection with correct direction between the continuous parents and the binary child. These results clearly suggest an enhanced performance of `DAGBagM` over both `DAGBagM`_*C*_ and `bnlearnD` in terms of directed edges detection when the binary node is the child.

**Table 3:** **Simulation scenario (ii)** – mixture of continuous and binary nodes. True DAG: *p* = 11, 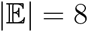; Sample size *n* = 50, 75, 100, SNR ∈ [0.5, 1.5]

| Method | $n$ | power | FDR | $F_1$ score | Percentage of correct edge detection from continuous parents to the binary child | |
| --- | --- | --- | --- | --- | --- | --- |
|  |  |  |  |  | both edges | at least one edge |
| DAGBagM | 50 | 0.89 | 0.1 | 0.89 | 0.34 | 0.84 |
| DAGBagM<sub>C</sub> |  | 0.87 | 0.1 | 0.88 | 0.15 | 0.67 |
| bnlearnD |  | 0.58 | 0.19 | 0.68 | 0.0 | 0.36 |
| DAGBagM | 75 | 0.95 | 0.09 | 0.93 | 0.57 | 0.92 |
| DAGBagM<sub>C</sub> |  | 0.9 | 0.1 | 0.90 | 0.17 | 0.62 |
| bnlearnD |  | 0.7 | 0.13 | 0.78 | 0.02 | 0.30 |
| DAGBagM | 100 | 0.97 | 0.08 | 0.94 | 0.78 | 0.98 |
| DAGBagM<sub>C</sub> |  | 0.92 | 0.08 | 0.92 | 0.31 | 0.75 |
| bnlearnD |  | 0.76 | 0.1 | 0.82 | 0.06 | 0.36 |

**Table 4:**
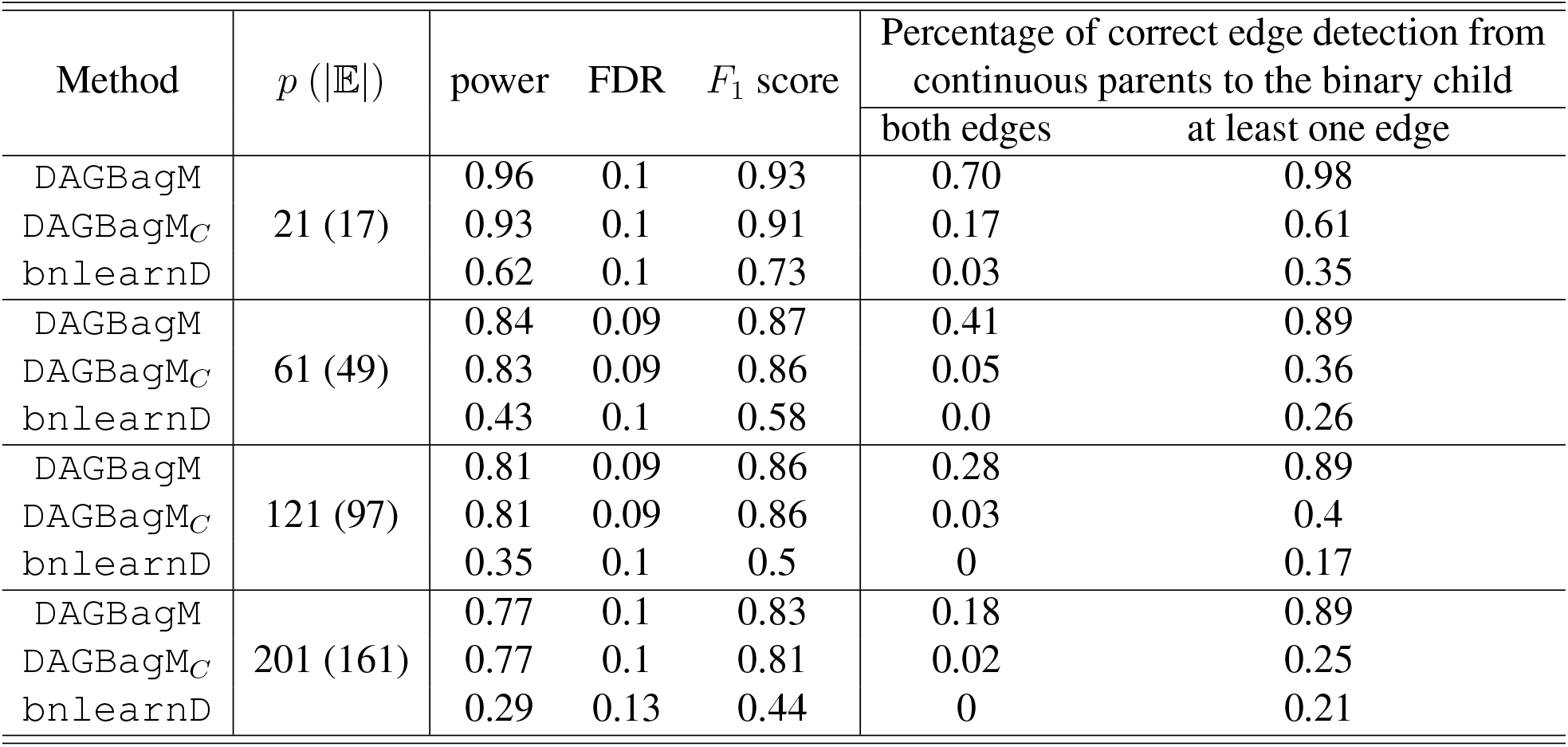
**Simulation scenario (ii)** – mixture of continuous and binary nodes with various combinations of *p* (number of nodes) and 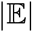 (number of edges) in the true DAG. Sample size *n* = 100, SNR ∈ [0.5, 1.5]

| Method | $p$ ($\vert \mathbb{E}\vert $) | power | FDR | $F_1$ score | Percentage of correct edge detection from continuous parents to the binary child | |
| --- | --- | --- | --- | --- | --- | --- |
|  |  |  |  |  | both edges | at least one edge |
| DAGBagM | 21 (17) | 0.96 | 0.1 | 0.93 | 0.70 | 0.98 |
| DAGBagM<sub>C</sub> |  | 0.93 | 0.1 | 0.91 | 0.17 | 0.61 |
| bnlearnD |  | 0.62 | 0.1 | 0.73 | 0.03 | 0.35 |
| DAGBagM | 61 (49) | 0.84 | 0.09 | 0.87 | 0.41 | 0.89 |
| DAGBagM<sub>C</sub> |  | 0.83 | 0.09 | 0.86 | 0.05 | 0.36 |
| bnlearnD |  | 0.43 | 0.1 | 0.58 | 0.0 | 0.26 |
| DAGBagM | 121 (97) | 0.81 | 0.09 | 0.86 | 0.28 | 0.89 |
| DAGBagM<sub>C</sub> |  | 0.81 | 0.09 | 0.86 | 0.03 | 0.4 |
| bnlearnD |  | 0.35 | 0.1 | 0.5 | 0 | 0.17 |
| DAGBagM | 201 (161) | 0.77 | 0.1 | 0.83 | 0.18 | 0.89 |
| DAGBagM<sub>C</sub> |  | 0.77 | 0.1 | 0.81 | 0.02 | 0.25 |
| bnlearnD |  | 0.29 | 0.13 | 0.44 | 0 | 0.21 |

The simulation results suggest that aggregation is an effective way to reduce false positives in DAG structure learning and treating continuous nodes and binary nodes using different models are beneficial in presence of mixture types of nodes in terms of both edge and edge direction detection.

We also compare the run time of the HC algorithm implemented in `DAGBagM` with the HC implementation in the R package `bnlearn`. Figure 3 shows the run time as a function of the number of nodes (edges) with sample size fixed at *n* = 500 and maximum number of search steps capped at 1000. As can be seen from Figure 3 the run time of `HC_bnlearn` increases dramatically as the number of nodes increases, while the runtime of `HC_DAGBagM` increases at a much slower rate.

### 3.2 Application to ovarian cancer

Comprehensive *mass-spectrometry* based proteomics characterization have been carried out in multiple recent cancer studies (Zhang et al., 2016). Pathway activities characterized by proteomics data revealed surprising new information of tumor samples. Specifically, metabolic pathways such as *Oxidative Phosphorylation* and *Adipogenesis* are the ones showing the least correlation between RNA and proteomics data in multiple tumors, suggesting active post-translational modifications to the members of these pathways in tumors. Metabolic reprogramming, recognized as one of the cancer hallmarks (Phan et al., 2014), promotes the activation of oncogenes and thus facilitates cancer progression and metastasis (Boroughs and DeBerardinis, 2016). This motivates us to screen for potential prognostic protein markers in related pathways in ovarian cancer based on newly generated proteomics data, which might lead to new insights missed in previous genomic based studies. Specifically, we apply `DAGBagM` on multiple ovarian cancer proteomics data sets, focusing on Oxidative Phosphorylation and Adipogenesis pathways, to derive a causal graph to characterize the dependence of patient progression on protein marker activities.

The detailed data analysis pipeline is illustrated in Figure 2 and Supplementary Table in Section A.3. Briefly, the pipeline consists of three major steps. We first derive prior information on causal protein-protein interactions using a time-course ovarian cancer cell line data set (step 1). Then, based on the direction information learned from step 1, we construct a DAG for proteins in the Oxidative Phosphorylation and Adipogenesis pathways based on a tumor proteomics data set from the CPTAC prospective ovarian (*Prosp-ova*) cancer study (step 2). We then extract closely linked network modules (small subsets of proteins) from the inferred DAG in step 2. For each module, we construct an outcome-driven-DAG based on an independent tumor proteomics data set from the CPTAC-TCGA retrospective ovarian (*Retro-Ova*) cancer study where patient survival outcome information is available (Zhang et al., 2016) (step 3). Throughout, we focus on the 260 proteins from the Oxidative Phosphorylation and Adipogenesis pathways that were observed in all three proteomics data sets. Here instead of learning the prognosis networks using the *Retro-Ova* data directly, we perform the intermediate step 2 in order to leverage the larger sample size of the *Prosp-ova* data and to confine the prognostic networks learning to small modules identified in step 2.

**Figure 2:**
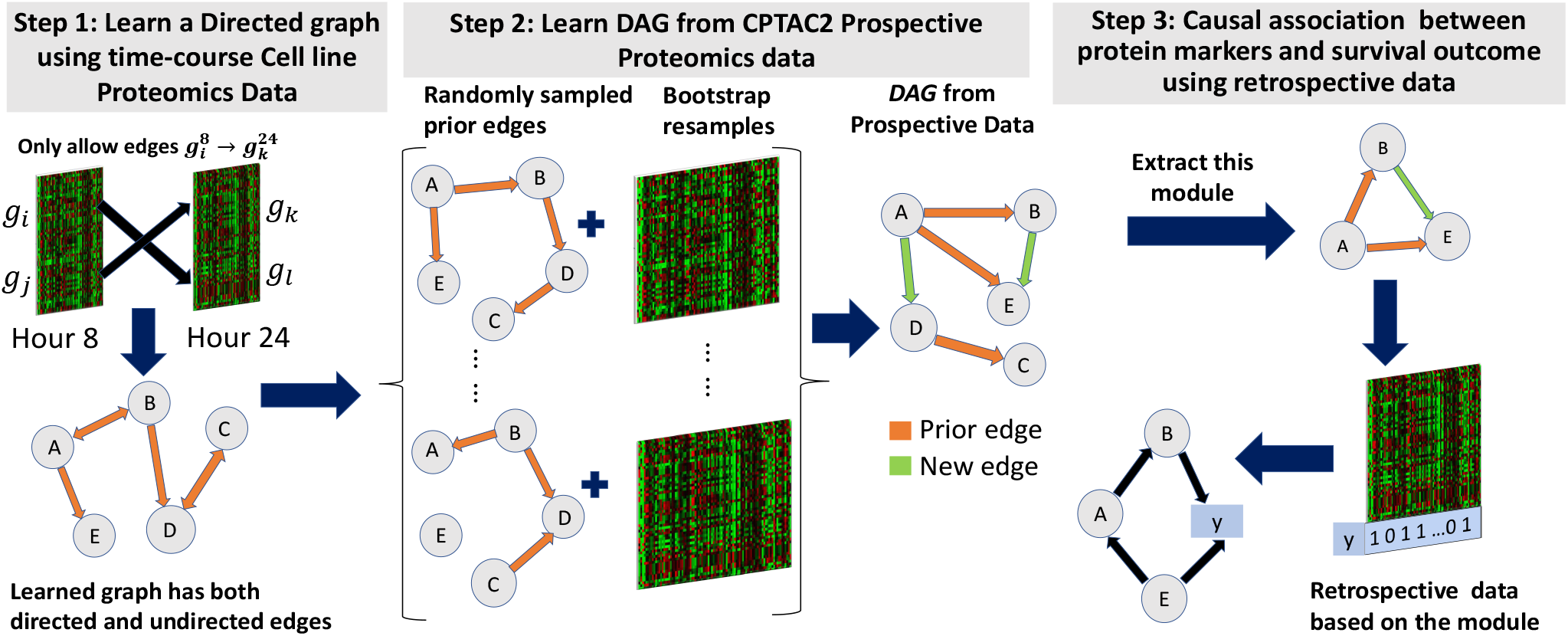
Application: integrative DAG learning analysis pipeline. In step 1 we obtain a directed network from cell line proteomics data, in step 2 we learn a protein DAG from CPTAC2 prospective data based on the direction information learned in step 1, and finally in step 3 we learn outcome-driven-DAGs with both continuous (protein markers) and binary variables (clinical outcome) based on the modules extracted from the DAG learned in step 2.

**Figure 3:**
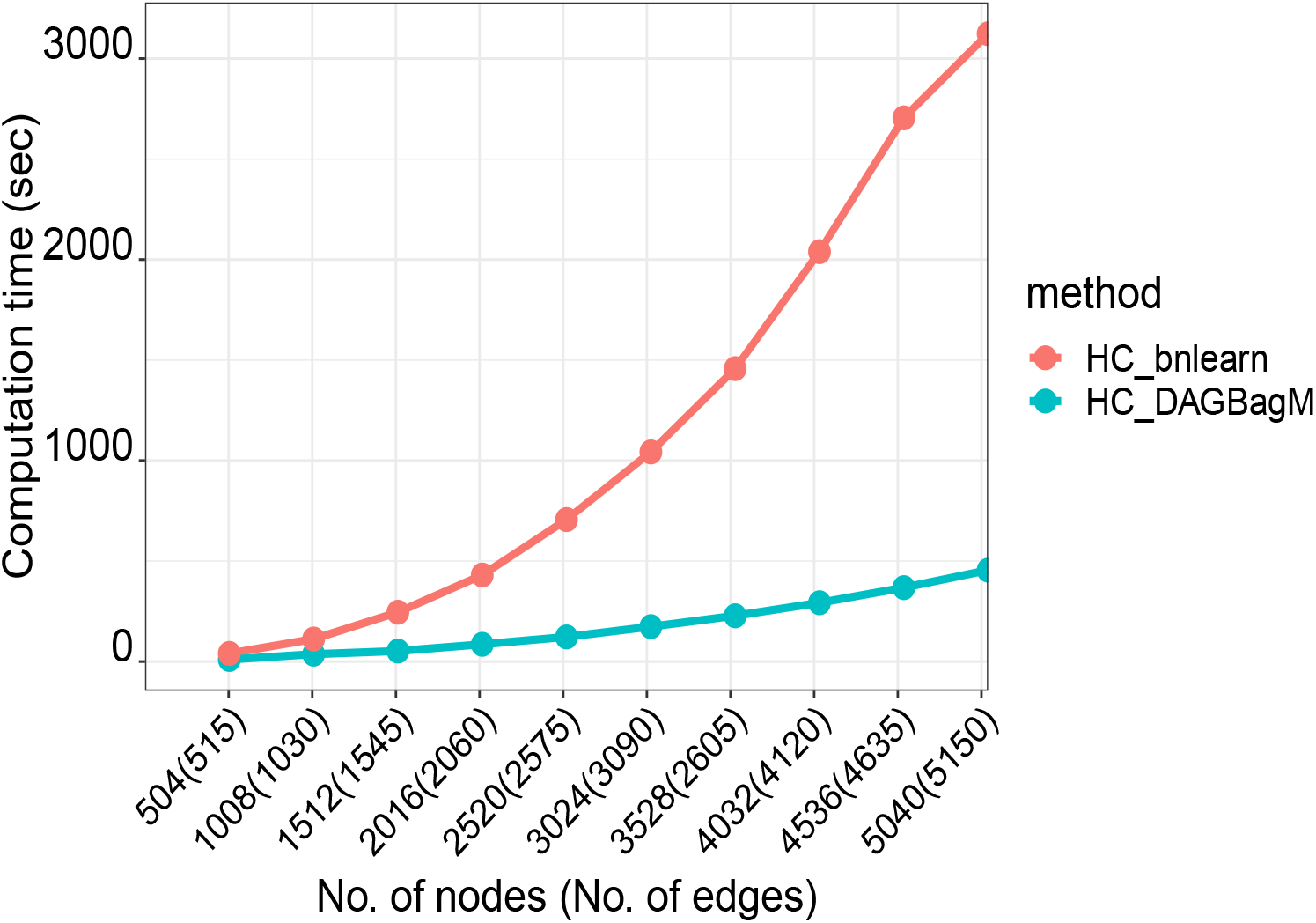
Run time comparison between HC implemented in `DAGBagM` (`HC_DAGBagM`) vs. HC implemented in `bnlearn` (`HC_bnlearn`) for different combination of nodes and edges.

#### Step 1: Learning regulatory direction on time-course cell line data

We first use a time-course cell line proteomics data (paper under preparation) to generate initial information on regulatory directions among the 260 proteins. The data contains proteomics profiles of 6 ovarian cancer cell lines, with three biological replicates of each cell line, from two different time points, 8 hours and 24 hours, after a chemo-drug perturbation. There are 36 proteomics profiles in total. It is reasonable to assume that the activities at the earlier time point drive the responses at the latter time point. Thus, for each protein, we treat its measurements at the two time points as two separate nodes and we create a blacklist that excludes edges from a node at the 24-hour time point to a node at the 8-hour time point. By applying the `DAGBagM` algorithm, we identify 100 directed edges among the 520 nodes. Out of these nodes, 81 nodes have at least one child and 100 nodes have at least one parent. Due to the small sample size, we can expect that only part of the regulatory relationships is identified in the estimated DAG. However, this provides valuable prior information for the step 2 analysis based on a larger data set.

#### Step 2: Identify well connected network modules based on Prosp-Ova data

Next, we learn a DAG using the Prosp-Ova proteomics data (McDermott et al., 2020; Arshad et al., 2019) of *n* = 108 treatment naive primary tumor samples, leveraging the direction information learned from the cell line data (step 1) as prior information. The Prosp-Ova proteomics data is obtained from CPTAC data portal (https://cptac-data-portal.georgetown.edu/cptac/s/S039) and data preporcessing is described in A.5.1 of the Supplement Material. We incorporate the prior information by specifying a whitelist while learning each individual DAG on a bootstrap resample. More specifically, since some of the directions learned from the cell line data could be either false positives or do not apply to tumor cells, along with each bootstrap resample of the data, we also randomly sample a subset of the inferred edges from step 1 to form a whitelist. By this way, false edges from step 1 will only impact a subset of the DAGs in the ensemble and thus are more likely to be filtered out during the aggregation step.

The resulting DAG contains 188 directed edges among the 260 proteins (Fig. 4A). Specifically, 113 proteins have at least one child node and 160 have at least one parent node. This DAG reveals several well connected network modules, each containing roughly 10-20 proteins after we apply the “cluster edge betweenness” function implemented in the R package *igraph* (Csardi and Nepusz, 2006). We then derive outcome driven DAGs for each of these modules based on the Retro-Ova data as described in step 3.

**Figure 4:**
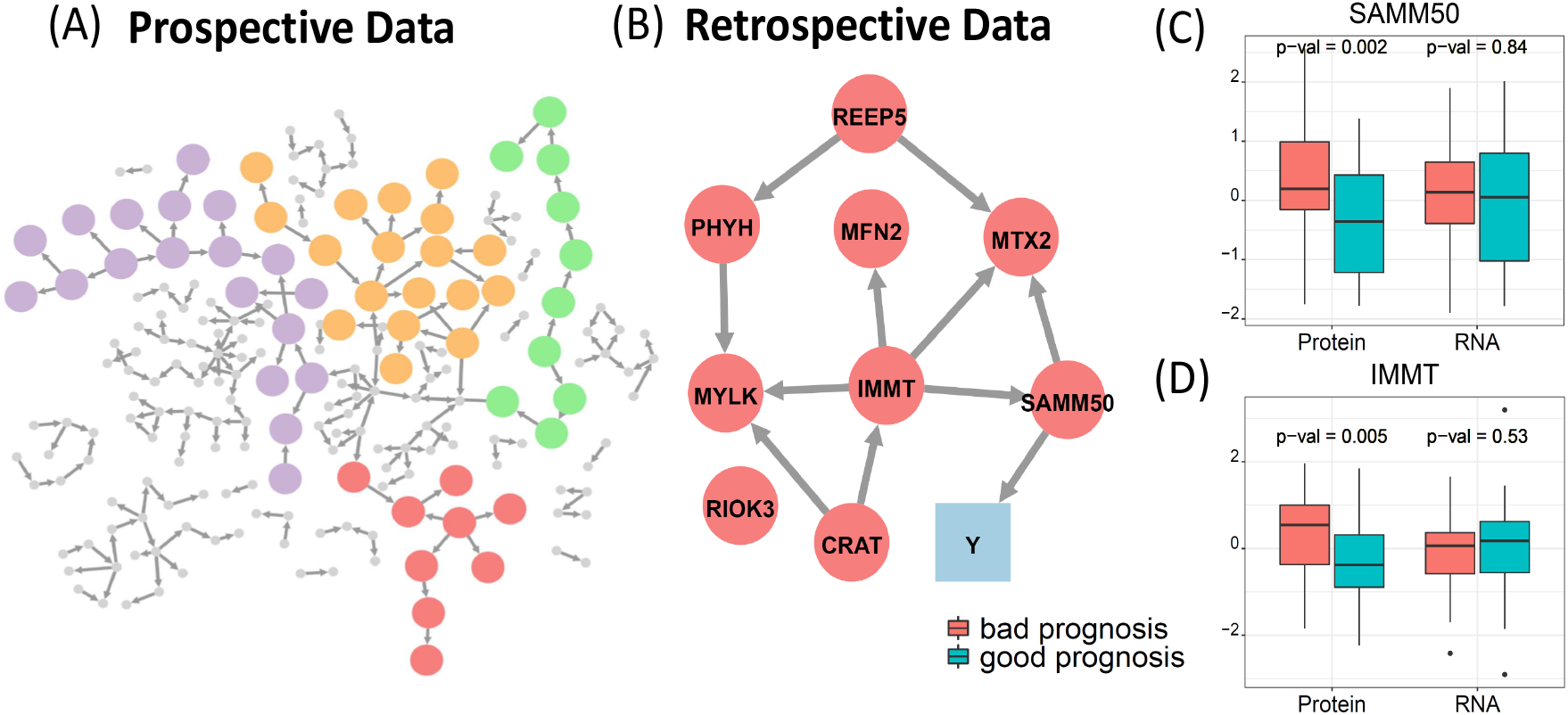
Application to ovarian cancer. (A) shows DAG learned from the Prosp-Ova data; (B) shows DAG learned from Retro-Ova data on the red module highlighted in (A); (C) and (D) respectively shows the boxplot of protein abundance and RNA expression level of SAMM50 and IMMT in the good and bad prognostic groups based on Retro-Ova data.

#### Step 3: Identifying prognostic protein biomarkers using the Retro-Ova data

In the final step of the integrative DAG learning, we seek for potential prognostic protein markers of ovarian cancer based on the Retro-ova data, which contains proteomics profiles of treatment naive primary tumor samples and *overall survival* (*OS*) information of 174 ovarian cancer patients (Zhang et al., 2016). Since OS of cancer patients is a very noisy outcome, depending on many factors in addition to the tumor characteristics, we focus on a subgroup of patients with extreme OS to enhance the power of detecting protein signals relevant to tumor progression. Specifically, we define good and bad prognosis status using *OS* > 5.5 years and *OS* < 1.5 years, respectively. There are 43 patients classified to the good prognosis group; and 36 to the bad prognosis group. Then for each module from step 2, we apply `DAGBagM` to the abundance measures of proteins in the module together with a binary variable *Y* representing the prognosis status across the 79 selected patients to seek for prognostic protein biomarkers. Figure 4 and Supplementary Figure A.3 illustrate the inferred DAGs of four modules in which causal protein markers for prognostic status were suggested.

The estimated DAG of the Red Module (Fig. 4B) is of particular interest, in which SAMM50 is a parent node causally associated with the prognosis status, with an upstream regulator, IMMT. For both SAMM50 and IMMT, significantly higher protein abundances were observed in tumors with bad prognosis than those with good prognosis (Fig. 4C and 4D). Protein of SAMM50 is a component of the sorting and assembly machinery of the mitochondrial outer membrane. It was hypothesized that change in the transport of proteins and metabolites into mitochondria due to SAMM50 is responsible for the energy production in cancer cells, and thus activity of SAMM50 has been suggested to be predictive of cancer progression in breast cancer(Sharma, 2011). SAMM50 protein closely associates with the mitochondrial contact site and cristae organizing system (MICOS) complex (Ott et al., 2015), of which IMMT is an important player. Recently, study has revealed the prognostic value of IMMT protein in gastric cancer (Sotgia and Lisanti, 2017). Intriguingly, the above result of `DAGBagM` analysis, for the first time, suggests the similar prognostic roles of proteins of SAMM50 and IMMT in ovarian cancer.

On the other hand, based on RNAseq data of the same set of tumors (Zhang et al., 2016), no significant association were detected between RNA expression levels of either SAMM50 or IMMT with prognositc status (Fig. 4C and 4D). Since proteins of SAMM50 and IMMT relate to the MICOS protein complex, likely their protein abundances are greatly influenced by post-translational modifications during complex forming and/or activation. Indeed, RNA expression level and protein abundance of both SAMM50 and IMMT in the same tumor samples showed poor correlation across patients (Supplementary Figure A.4). These findings nicely illustrate the unique potential of proteomics data based investigation.

We also applied `bnlearnD` on the same module, where IMMT and MYLK were inferred as parent nodes of the prognosis status (Supplementary Figure A.5 A). We observe that there is no significant difference in protein abundance of MYLK between the tumors with good and bad prognosis status (Supplementary Figure A.5 B). This indicates that the edge between MYLK and the prognosis status inferred by `bnlearnD` is possibly a false positive.

## 4 Discussion

In this paper, we propose `DAGBagM`, a novel DAG structure learning tool for data with both continuous and binary variables using a score-based method coupled with bootstrap aggregation. Our contributions are threefold. First, we propose a score-based DAG structure learning algorithm which allows for (i) both continuous and binary nodes; and (ii) either type of nodes being a child node. As shown by simulation studies, `DAGBagM` achieves better performance for detecting edges and edge directions, compared to conventional methods and strategies which either do not allow binary children or treat all nodes as one type. Such flexibility provided by `DAGBagM` has important relevance in practice when one is interested in detecting biomarkers causally associated with clinical outcomes, as the latter are often binary endpoints and the former are often continuous measurements. Second, we propose a novel technique to aggregate DAGs learned on bootstrap resamples using a structural Hamming distance on the DAG space. Simulation results show that the proposed aggregation strategy is able to greatly reduce the number of false positives. Moreover, this aggregation procedure is a general tool that can be coupled with any structure learning algorithm (score based or not) and is a flexible way to incorporate prior information. Lastly, we develop an efficient implementation of the hill-climbing algorithm by utilizing information from previous search steps which greatly speeds up both score calculation and acyclic status checking. Particularly, our implementation is much faster than that in a widely used DAG learning R package `bnlearn`.

Based on `DAGBagM`, we then implemented an integrative DAG learning pipeline to analyze multiple ovarian cancer proteomics datasets with the goal to identify potential prognostic proteins in ovarian cancer. In DAG structure learning, it is very challenging to identify edge directions unambiguously due to identifiability issues. To facilitate the inference of edge directions, we utilize a time-course cell line proteomics data to get the initial regulatory direction estimation. We then use learned edges as prior information to construct DAGs from two larger tumor proteomics data sets. Specifically, we utilize the Prosp-Ova data, which does not contain patient prognosis information, for dimension reduction by deriving network modules based on its inferred DAG; while utilize the Retro-Ova data to derive the final outcome-driven-DAG of each network module. Such a strategy increases the power of identifying meaningful causal relationships.

Our result reveals multiple candidate protein biomarkers, including SAMM50 and IMMT, to be causally associated with cancer prognosis. Proteins of SAMM50 and IMMT are important regulator and member of the MICOS complex, respectively. While prognostic values of SAMM50 and IMMT have been reported in other cancer types, our analysis for the first time suggests their prognostic roles in ovarian cancer. Intriguingly, the prognostic value of SAMM50 and IMMT could not be observed based on RNAseq data from the same set of patients, suggesting the importance and great potential of employing proteogenomic integrative analysis in biomedical research. The markers identified to have a causal relationship with the survival outcome in ovarian cancer dataset could serve as potential targets to individualized anti-cancer agents, upon evaluation through clinical practice (Kamel and Al-Amoudi, 2017). In this paper, we focus on the Oxidative Phosphorylation and Adipogenesis pathways to investigate the underlying molecular mechanism of their members. Although extending the analysis pipeline developed here to other pathways in a systematic manner is out of the scope of this paper, we see great promises of `DAGBagM` in systems biology applications.

## Acknowledgements

We would like to thank Drs. Karin Rodland, Hui Zhang and their teams to share the ovarian proteomics data from the National Cancer Institutes Clinical Proteomic Tumor Analysis Consortium (CPTAC) studies.

