## Supplementary file for "DAGBagM: Learning directed acyclic graphs of mixed variables with an application to identify prognostic protein biomarkers in ovarian cancer"

### SUPPLEMENTARY MATERIAL

#### A.1 Efficient implementation of hill climbing algorithm

In the following, we use $\tilde{\mathcal{G}}$ to denote the graph in the previous step, $O^*$ to denote the selected operation in the previous step, and $\mathcal{G}^* = O^*(\tilde{\mathcal{G}})$ to denote the current graph. We also use $\delta(O : \mathcal{G})$ to denote the change of score resulting from applying operation $O$ to graph $\mathcal{G}$. We summarize the score updating and acyclic check schemes in the following two propositions.

**Proposition 1** *Suppose score$(\cdot : \cdot)$ is a decomposable score. For an (eligible) operation $O$, $\delta(O : \tilde{\mathcal{G}}) = \delta(O : \mathcal{G}^*)$, if one of the following holds:*

- *$O^*$ is one of the forms: "add $x_i \to x_j$" or "delete $x_i \to x_j$", and $O$ is not one of the forms: "add $x_k \to x_j$" , "delete $x_k \to x_j$" , " reverse $x_j \to x_k$ ".*
- *$O^*$ is of the form "reverse $x_i \to x_j$", and $O$ is not one of the forms: "add $x_k \to x_j$" , "delete $x_k \to x_j$" , " reverse $x_j \to x_k$ ", "add $x_k \to x_i$" , "delete $x_k \to x_i$", " reverse $x_i \to x_k$ ".*

In short, any operation that does not involve the neighborhoods changed by the selected operation in the previous step will lead to the same score change as in the previous step. This has been pointed out by Koller and Friedman (2009).

In the following, $de(x_i)$ denotes the set of descendant of node $x_i$ and $an(x_i)$ denotes the set of ancestors of node $x_i$.

**Proposition 2** *The following holds for acyclic check.*

- *If $O^*$ is of the form "add $x_{i^*} \to x_{j^*}$", then for an operation $O$ the following holds:*
  - *If $O$ does not lead to cycles in the previous step, then*
    - *if $O$ is of the form "add $x_i \to x_j$", and $i \in de(x_{j^*})$ and $j \in an(x_{i^*})$, then $O$ leads to a cycle.*
    - *if $O$ is of the form "reverse $x_i \to x_j$", and $j \in de(x_{j^*})$ and $i \in an(x_{i^*})$, then $O$ leads to a cycle.*
    - *if otherwise, $O$ remains acyclic.*
  - *If $O$ leads to a cycle in the previous step, it remains cyclic.*
- *If $O^*$ is of the form "delete $x_{i^*} \to x_{j^*}$", then for an operation $O$ the following holds:*
  - *If $O$ does not lead to cycles in the previous step, it remains acyclic.*
  - *If $O$ leads to a cycle in the previous step, then*
    - *if $O$ is of the form "add $x_i \to x_j$", and $i \in de(x_{j^*})$ and $j \in an(x_{i^*})$, then we need to check its acyclicity;*
    - *if $O$ is of the form "reverse $x_i \to x_j$", and $j \in de(x_{j^*})$ and $i \in an(x_{i^*})$, then we need to check its acyclicity;*
    - *if otherwise, $O$ remains cyclic.*
- *If $O^*$ is of the form "reverse $x_{i^*} \to x_{j^*}$", then for an operation $O$ the following holds:*

- *If $O$ does not lead to cycles in the previous step, then*
  - *if $O$ is of the form "add $x_i \to x_j$", and $i \in de(x_{i^*})$ and $j \in an(x_{j^*})$, then $O$ leads to a cycle.*
  - *if $O$ is of the form "reverse $x_i \to x_j$", and $j \in de(x_{i^*})$ and $i \in an(x_{j^*})$, then $O$ leads to a cycle.*
  - *if otherwise, $O$ remains acyclic.*
- *If $O$ leads to a cycle in the previous step, then*
  - *if $O$ is of the form "add $x_i \to x_j$", and $i \in de(x_{j^*})$ and $j \in an(x_{i^*})$, then we need to check its acyclicity;*
  - *if $O$ is of the form "reverse $x_i \to x_j$", and $j \in de(x_{j^*})$ and $i \in an(x_{i^*})$, then we need to check its acyclicity;*
  - *if otherwise, $O$ remains cyclic.*

#### A.2 Bootstrap Aggregation using hill climbing algorithm

Given an ensemble of DAGs: $\mathbb{G}^e = \{\mathcal{G}_b : b = 1, \cdots B\}$, the selection frequency (SF) of a directed edge $e$, when the reversal of an edge is counted as one unit of operation, is defined as

$$gp_e := p_e + \frac{1}{2} p_{e^*},$$

where $e^*$ denotes the edge with the reversed direction of $e$. $score_d$ can be expressed in terms of SF as given in the following proposition:

**Proposition 3** *Given an ensemble of DAGs:* $\mathbb{G}^e = \{\mathcal{G}_b : b = 1, \cdots B\}$*, the aggregation score under* $d$ *is*

$$score_d(\mathcal{G} : \mathbb{G}^e) = \sum_{e \in \mathbb{E}(\mathcal{G})} (1 - 2gp_e) + C,$$

*where*

$$C = \frac{1}{B} \sum_{b=1}^{B} \sum_{i=1}^{p} \sum_{j=1}^{p} \mathbb{A}_b(i,j) = \sum_{i=1}^{p} \sum_{j=1}^{p} p_{x_i \to x_j},$$

*is a constant which only depends on the ensemble* $\mathbb{G}^e$*, but does not depend on* $\mathcal{G}$*.*

**Proof of Proposition 3**. To help with the proof, we use Table A.1 to show the value of $S_{ij}(\mathbb{A}, \tilde{\mathbb{A}}) = \max\{|\mathbb{A}(i,j) - \tilde{\mathbb{A}}(i,j)|$, for $1 \leq i < j \leq p$. For an adjacency matrix $\mathbb{A}$ and a given pair $(i,j)$ with $i < j$, let $(1,0)$ denote the case where $\mathbb{A}(i,j) = 1, \mathbb{A}(j,i) = 0$, $(0,1)$ denote the case where $\mathbb{A}(i,j) = 0, \mathbb{A}(j,i) = 1$ and $(0,0)$ denote the case where $\mathbb{A}(i,j) = 0, \mathbb{A}(j,i) = 0$ (note $(1,1)$ is not possible due to the acyclic constraint).

Table A.1: $S_{ij}(\mathbb{A}, \tilde{\mathbb{A}})$ for $1 \le i < j \le p$.

| $\tilde{\mathbb{A}} \backslash \mathbb{A}$ | $(1,0)$ | $(0,1)$ | $(0,0)$ |
|---|---|---|---|
| $(1,0)$ | 0 | 1 | 1 |
| $(0,1)$ | 1 | 0 | 1 |
| $(0,0)$ | 1 | 1 | 0 |

By definition:

$$
\begin{aligned}
score_d(\mathcal{G} : \mathbb{G}^e) &= \frac{1}{B} \sum_{b=1}^{B} \sum_{1 \le i < j \le p} S_{ij}(\mathbb{A}, \mathbb{A}_b) \\
&= \sum_{i<j: x_i \to x_j \in \mathbb{E}(\mathcal{G})} \left(p_{ji} + p_{ij}^0\right) \\
&+ \sum_{i<j: x_j \to x_i \in \mathbb{E}(\mathcal{G})} \left(p_{ij} + p_{ij}^0\right) \\
&+ \sum_{i<j: x_i \to x_j \notin \mathbb{E}(\mathcal{G}), x_j \to x_i \notin \mathbb{E}(\mathcal{G})} \left(p_{ij} + p_{ji}\right),
\end{aligned}
$$

where $p_{ij}$ denotes the selection frequency of edge $x_i \to x_j$, and $p_{ij}^0 = p_{ji}^0 := 1 - p_{ij} - p_{ji}$ is the frequency that neither $x_i \to x_j$ nor $x_j \to x_i$ got selected.

Note that,

$$
\begin{aligned}
\sum_{i<j: x_i \to x_j \notin \mathbb{E}(\mathcal{G}), x_j \to x_i \notin \mathbb{E}(\mathcal{G})} \left(p_{ij} + p_{ji}\right) &= \sum_{1 \le i < j \le p} \left(p_{ij} + p_{ji}\right) \\
&- \sum_{i<j: x_i \to x_j \in \mathbb{E}(\mathcal{G})} \left(p_{ij} + p_{ji}\right) - \sum_{i<j: x_j \to x_i \in \mathbb{E}(\mathcal{G})} \left(p_{ij} + p_{ji}\right).
\end{aligned}
$$

Therefore

$$\begin{aligned} score_d(\mathcal{G} : \mathbb{G}^e) &= \sum_{i<j: x_i \to x_j \in \mathbb{E}(\mathcal{G})} (p_{ji} + 1 - 2(p_{ij} + p_{ji})) \\ &+ \sum_{i<j: x_j \to x_i \in \mathbb{E}(\mathcal{G})} (p_{ij} + 1 - 2(p_{ij} + p_{ji})) \\ &+ \sum_{1 \le i<j \le p} (p_{ij} + p_{ji}) \\ &= \sum_{i<j: x_i \to x_j \in \mathbb{E}(\mathcal{G})} \left(1 - 2\left(p_{ij} + \frac{1}{2}p_{ji}\right)\right) \\ &+ \sum_{i<j: x_j \to x_i \in \mathbb{E}(\mathcal{G})} \left(1 - 2\left(p_{ji} + \frac{1}{2}p_{ij}\right)\right) \\ &+ \sum_{1 \le i<j \le p} (p_{ij} + p_{ji}). \end{aligned}$$

By definitions of the constant $C$ and the selection frequency, we complete the proof.

Also the hill climbing search algorithm with $score_d$ can be simplified to the procedure described in Table A.2.

Table A.2: Hill climbing algorithm for DAG aggregation

.

**Input:** an ensemble of DAGs: $\mathbb{G}^e = \{\mathcal{G}_b : b = 1, \cdots B\}$.

Calculate selection frequency (SF) $gp_e$ for all possible edges.

Order edges with SF > 50%.

Add edges sequentially according to SF and stop when SF $\leq 0.5$.

- Initial step: $\mathcal{G}^{(0)}$= empty graph, $\mathbb{C}$ = empty set.
- $s^{th}$ step: current graph $\mathcal{G}^{(s)}$, current operation $O$: "add the edge with the $s^{th}$ largest SF".
  - If $O$ passes the acyclic check, then $\mathcal{G}^{(s+1)} = O(\mathcal{G}^{(s)})$, i.e., add this edge. If $O$ does not pass the acyclic check, then $\mathcal{G}^{(s+1)} = \mathcal{G}^{(s)}$, i.e., does not add this edge, and add this edge to $\mathbb{C}$.
  - If the $(s+1)^{th}$ largest SF > 50%, proceed to step $s + 1$. Otherwise, set $\mathcal{G}^* = \mathcal{G}^{(s+1)}$ and stop the algorithm.

**Output:** $\mathcal{G}^*$, and the set of "cyclic edges" $\mathbb{C}$.

#### A.3 Integrative DAG learning pipeline for real data application

Integrative DAG learning pipeline for real data application

1: Consider the $n_1 = 18$ samples from the pre-processed cell line proteomics data $D_1$ perturbed by treatment at time-point $t_1 = 8hr$ and $n_2 = 18$ samples perturbed by treatment at time-point $t_2 = 24hr$.

2: Consider the $G = 260$ proteins from Adipogenesis and Oxidative Phosphorylation pathways that are observed in both $D_1$ and Prosp-ova data $D_2$: call it $M_G$, and denote each member of $M_G$ by $g_i$, $i = 1, \ldots, G$.

3: Stack $A$: matrix of proteins from $M_G$ by samples at $t_1$ with $B$: matrix with genes from $M_G$ by samples at $t_2$: call it $C_{(8+24)}$ which is a matrix of dimension $2G \times n_1$.

4: Apply `DAGBagM` to $C_{(8+24)}$ with blacklist $BL$ to learn a directed network from $D_1$: call it $C_{est}$ of dimension $2G \times 2G$. $BL$ is used to suppress all edges from (i) $g_i$ to $g_j$ at $t_1$, (ii) $g_i$ to $g_j$ at $t_2$, and (iii) $g_i$ at $t_2$ to $g_j$ at $t_1$, for all $(i, j)$.

5: Extract the top off-diagonal block matrix ($G \times G$) corresponding to the estimated edges directed from proteins at $t_1$ to proteins at $t_2$: call it $C_{est(8 \to 24)}$. Since there might be an edge from protein $g_i$ at $t_1$ to another protein $g_j$ at $t_2$ and another edge from $g_j$ at $t_1$ to $g_i$ at $t_2$, $C_{est(8 \to 24)}$ is not necessarily a DAG. In the next step, we select a subset of edges to pass as whitelist for learning a directed network from $D_2$.

6: Sample edges randomly to select 90% edges from $C_{est(8 \to 24)}$ and sequentially add them in an empty graph with a check of acyclic status at every step: call this network $C^*_{est(8 \to 24)}$ which is a DAG. Repeat this step 100 times and call each such DAG $C^*_{b\,est(8 \to 24)}$, $b = 1, \ldots, 100$

7: Generate $B = 100$ bootstrap resamples of the data $D_2$: call them $D_b$, $b = 1, \ldots, B$. Learn 100 DAGs, one from each of $D_b$ using $C^*_{b\,est(8 \to 24)}$ as prior (passed as whitelist). Aggregate 100 DAGs using our proposed aggregation procedure and call it $P_{est}$.

8: Identify small closely linked modules with $10 \sim 20$ proteins to from $P_{est}$.

9: Consider the Retro-ova data $D_3$ with $n_3 = 79$ samples of which $n_{3S}$ is index set of the patients with good survival and $n_{3R}$ is the index set of the patients with poor survival. Construct a vector $Y(1 \times n_3)$ whose elements are coded as 1/0 (1: $i \in n_{3S}$, 0: $i \in n_{3R}$) in $D_3$. Append $Y$ as an additional binary node to each extracted module.

10: Apply `DAGBagM` to learn a directed network from $D_3$ for each module.

#### A.4 Simulation: additional details

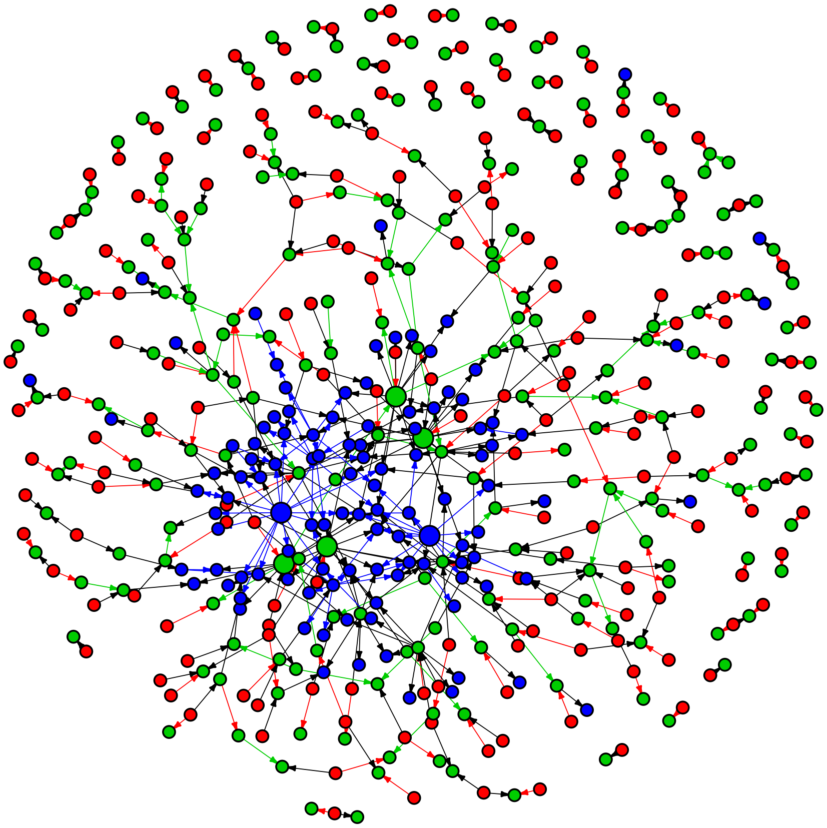

Figure A.1: Large Graph with $p = 504, |\mathbb{E}| = 515$.

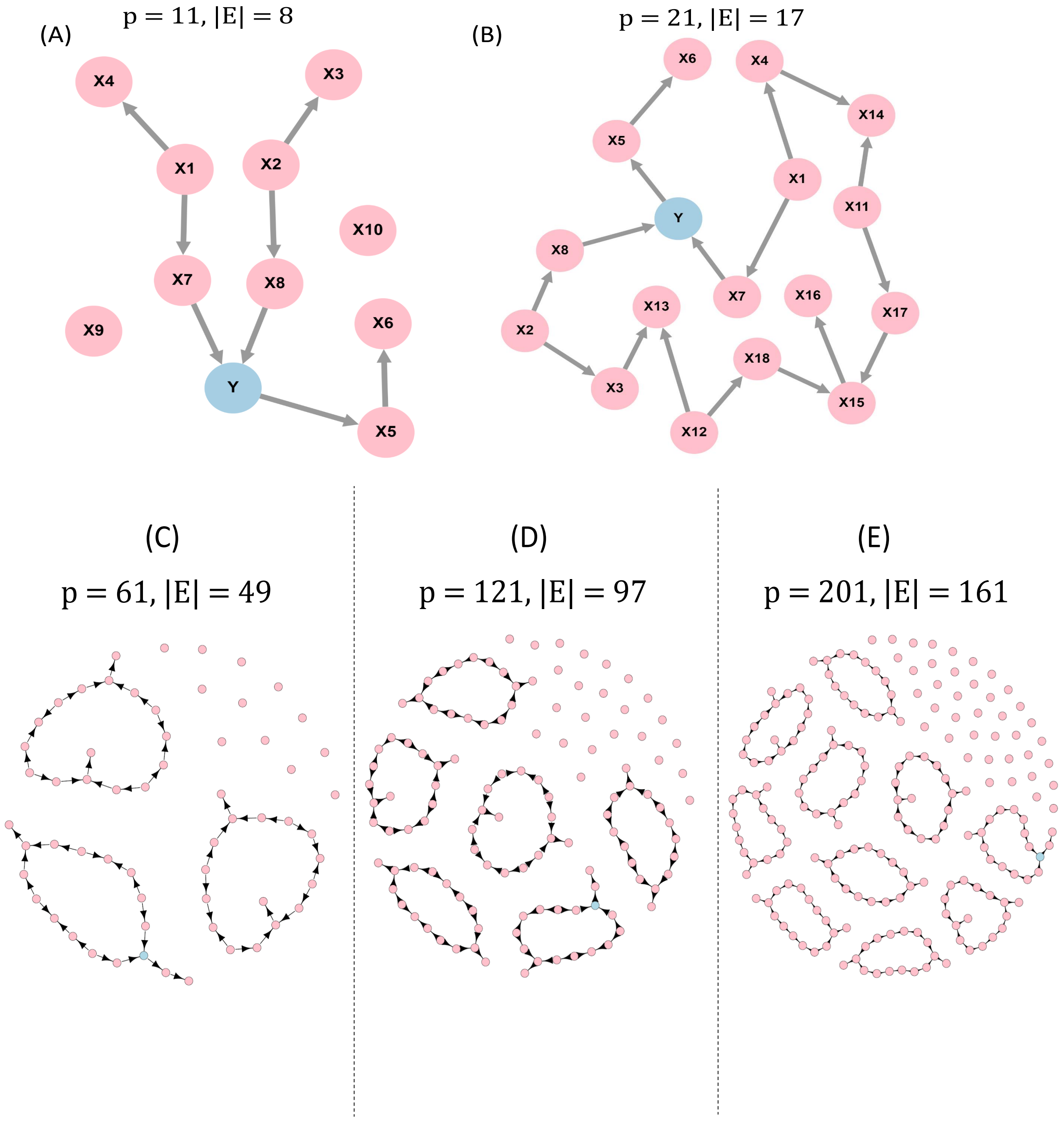


Figure A.2: (A) True DAG with $p = 11$, (10 continuous nodes and 1 binary node), $|\mathbb{E}| = 8$, (B) True DAG with $p = 21$, (20 continuous nodes and 1 binary node $Y$), $|\mathbb{E}| = 17$, (C) True DAG with $p = 61$, (60 continuous nodes and 1 binary node $Y$), $|\mathbb{E}| = 49$, (D) True DAG with $p = 121$, (120 continuous nodes and 1 binary node $Y$), $|\mathbb{E}| = 97$, (E) True DAG with $p = 201$, (200 continuous nodes and 1 binary node $Y$), $|\mathbb{E}| = 161$.

#### A.5 Application: Ovarian Cancer Proteomics Data

##### A.5.1 Pre-processing of the datasets used in three steps

We performed global normalization to align the sample median to remove any systematic variation across the samples for all the datasets. We then filtered out the proteins that are missing all samples in a given batch (batch-level missing). For Retro-ova and Prosp-ova data we again filtered out proteins that are missing from 75 % of the samples. We then applied batch correction to all three datasets using an R tool: *ComBat* (Johnson et al., 2007) to remove batch-effect. We also used an imputation tool *DreamAI* (https://github.com/WangLab-MSSM/DreamAI) to impute the Retro-ova and Prosp-ova data.

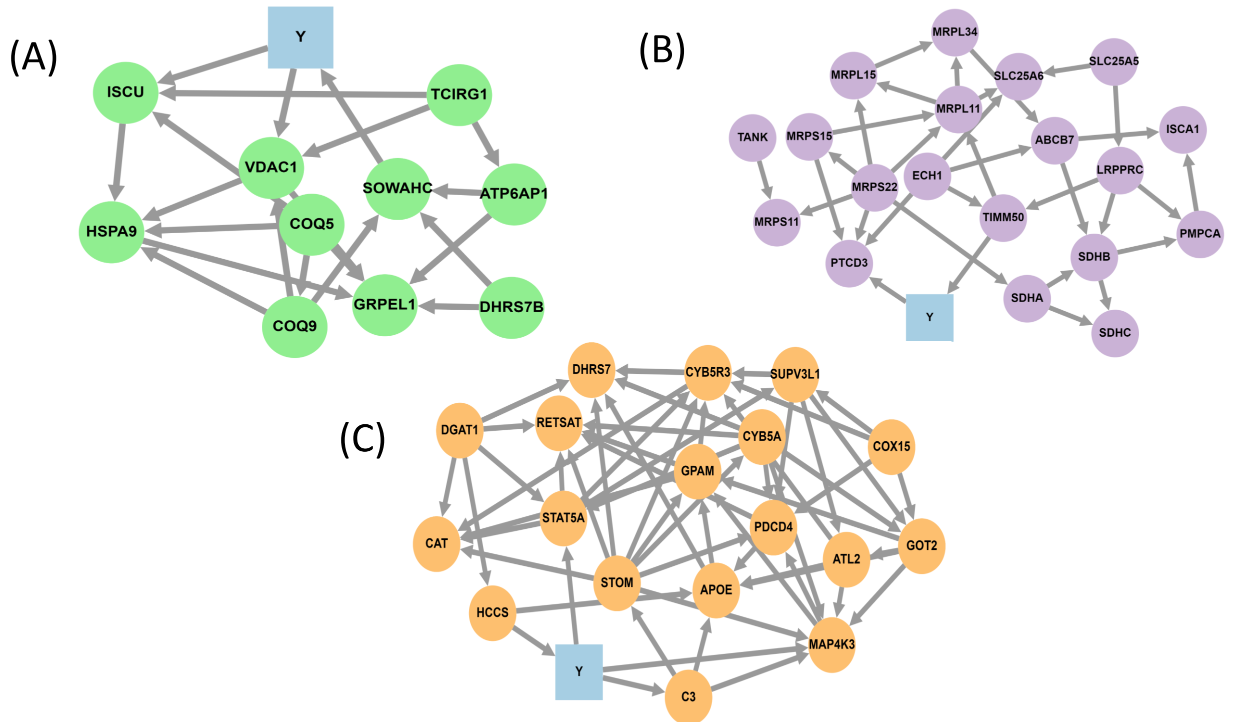


Figure A.3: DAGs based on (A) module 2 (green), (B) module 3 (purple) and (C) module 4 (orange) from Retro-ova data.

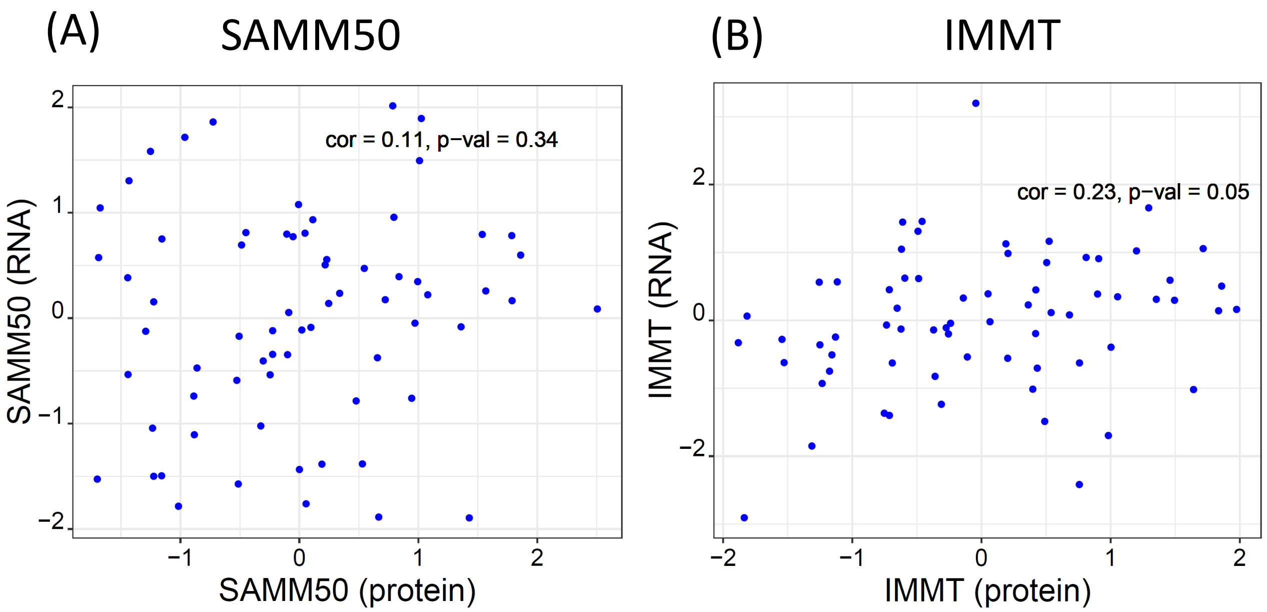


Figure A.4: (A) shows the scatterplot for the abundance of the protein SAMM50 from Retro-ova data vs. its RNA expression from CPTAC2, and (B) shows the scatterplot for the abundance of the protein IMMT from Retro-ova vs. its RNA expression from CPTAC2.

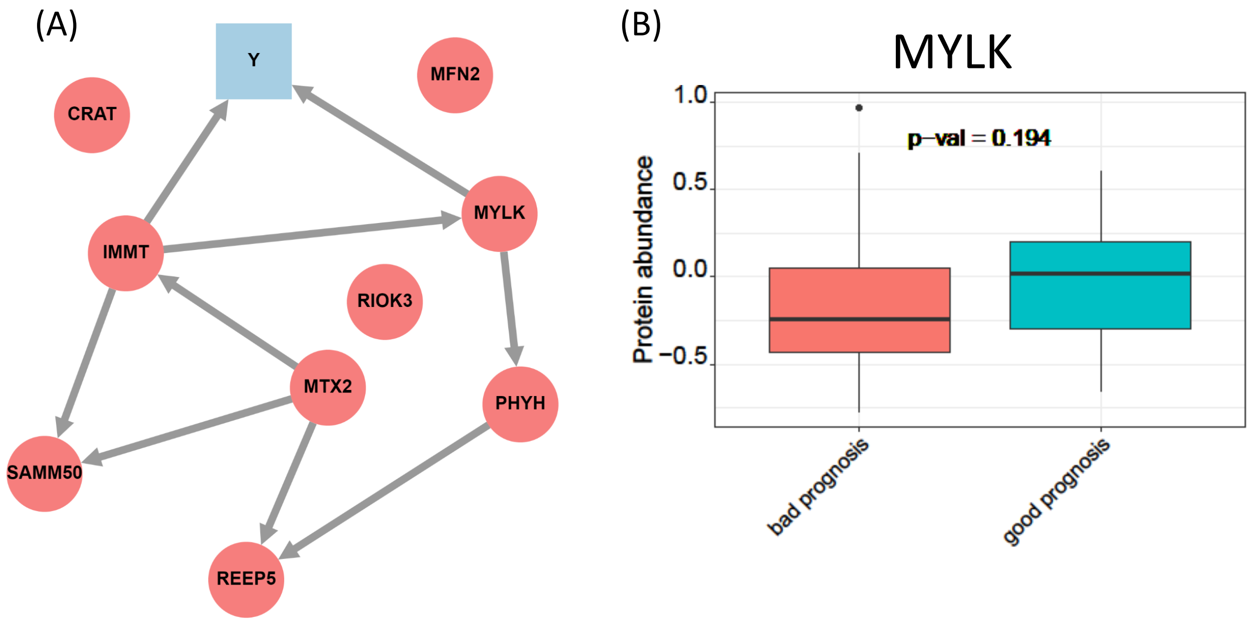


Figure A.5: (A) shows the estimated DAG based on module 1 (Red) by `bnlearnD`, and (B) shows the abundance of the protein MYLK based on Retro-ova data in the tumors with good and bad prognosis.
